# Kaiso regulates DNA methylation homeostasis

**DOI:** 10.1101/2020.12.09.417576

**Authors:** D Kaplun, G Filonova, Y. Lobanova, A Mazur, S Zhenilo

## Abstract

Gain and loss of DNA methylation in cells is a dynamic process that tends to achieve an equilibrium. Many factors are involved in maintaining the balance between DNA methylation and demethylation. Previously, it was shown that methyl-DNA protein Kaiso may attract NcoR, SMRT repressive complexes affecting histone modifications. On the other hand, the deficiency of Kaiso resulted in slightly reduced methylation of ICR in H19/Igf2 locus and Oct4 promoter in mouse embryonic fibroblasts. However, nothing is known whether Kaiso may attract DNA methyltransferase to influence DNA methylation level. The main idea of this work is that Kaiso may lead to DNA hypermethylation attracting de novo DNA methyltransferases. We demonstrated that Kaiso regulates TRIM25 promoter methylation. It can form a complex with DNMT3b. BTB/POZ domain of Kaiso and ADD domain of DNA methyltransferase are essential for complex formation. Thus, Kaiso can affect DNA methylation.

## INTRODUCTION

DNA methylation is crucial for organism development, cell differentiation, X inactivation, genomic imprinting, regulation of transcription. In vertebrates DNA methylation mainly occurs at CpG dinucleotides. Methylation of the promoter regions correlates with repression of transcription. About 70% of CpG are methylated except those that are a part of so called CpG islands. Establishment of DNA methylation is regulated by de novo DNA methyltransferases DNMT3a and DNMT3b. There are several mechanisms that are responsible for proper pattern of DNA methylation (for review (Greenberg and Bourc’his 2019)). Establishment of DNA methylation occurs during post-implantation period and changes during cell differentiation. The pattern of methylation remains relatively stable starting to change in aging period, during various pathological processes, after environmental impact. However, DNA methylation is dynamically changed in nervous cells during learning and memory formation (Day and Sweatt 2010; Oliveira 2016; Heyward and Sweatt 2015). It will be important to find new mechanisms involved in fast DNA methylation plasticity.

Deficiency of several methyl DNA proteins, MBD1, MBD2, MeCP2, Kaiso lead to behavioral deviations, changes in maternal behaviour or problems in learning and memory formation (Kulikov et al. 2016)(Ip, Mellios, and Sur 2018; Guy et al. 2001) (Hendrich et al. 2001), (Allan et al. 2008). It was hypothesized that the main function of these methyl DNA binding proteins lie in the area of memory and learning, but the mechanism of their involvement in DNA methylation dynamics is still undiscovered. One of these methyl DNA proteins Kaiso belongs to the BTB/POZ domain proteins family. It binds methylated DNA via three zinc fingers. Also, it can interact with sequences containing nonmethylated Kaiso binding site (KBS) CTGCNA, hydroxymethylaion prevents its binding to DNA (Prokhortchouk 2001; Daniel et al. 2002; S. V. Zhenilo, Musharova, and Pokhorchuk 2013; Zhigalova et al. 2015). Previously, it was shown that the deficiency of methyl-DNA binding protein Kaiso results in decreased methylation of ICR in H19/Igf2 imprinted loci and Oct4 promoter region ( (Bohne et al. 2016; Kaplun et al. 2019). Moreover, we demonstrated that transcriptional activity of Kaiso is dependent on posttranslational modification SUMOylation. DeSUMOylated Kaiso is able to completely inhibit TRIM25 promoter activity setting negative chromatin marks on it. TRIM25 promoter silence can not be reverted by presence of active SUMOylated form of Kaiso. The main goal of this work is to determine whether Kaiso can influence the establishment of DNA methylation and what is the mechanism underlying this. We demonstrated that deSUMOylated form of Kaiso not only established repressive histone marks but also involved in promoter TRIM25 hypermethylation. For the first time we showed that Kaiso can form a complex with DNMT3b de novo methyltransferase, but can not directly interact with it.

## MATERIALS AND METHODS

### Cell lines

HEK293, Kaiso KO and K42R Kaiso HEK293 cells (S. Zhenilo et al. 2018) were grown in Dulbecco’s modified Eagle medium supplemented with 10% fetal bovine serum, 1% penicillin/streptomycin, and 2 mm l-glutamine. Cells were transfected with Lipofectamine 2000 (ThermoFisherScientific) according to the manufacturer’s protocol. Cells were typically harvested 48 h post-transfection for further analysis.

### Plasmids and vectors

Kaiso-GFP, BTB-HA constructs were used the same as in (S. Zhenilo et al. 2018).

Full length Kaiso (1-692), BTB/POZ domain (1-117 amino acids), zinc fingers (494-573 amino acids), spacer (117-494 amino acids) were cloned in pGex-2T. pcDNA3/Myc-DNMT3B1 was a gift from Arthur Riggs (Addgene plasmid # 35522) (Chen, Mann, and Hsieh 2005). ADD, PWWP and catalytic domains of DNMT3b were amplified with primers ADDfor 5’-TTGAATTCGCACCCAAGCGCCTCAAGA and ADDrev 5’-TTGGATCCCCGTGTCACTGGTGAAGAAG, PWWPfor 5’-TTGAATTCGCAGACAGTGGAGATGGAGA-, PWWPrev 5’-TTGGATCCTCCAGAGCATGGTACATGG, catfor 5’-TTGAATTCCCTGCCATTCCCGCAGCC, catrev 5’-GCGGATCCTATTCACATGCAAA and subcloned to pcDNA3-myc vector.

### Antibodies

Following reagents were used in this study: anti-Kaiso polyclonal rabbit antibodies (kindly gifted by Dr. A. Reynolds), anti-HA (H6908, Sigma), anti-HA agarose (A2095, Sigma), anti-actin (ab8227), anti Kaiso 6F (ab12723, Abcam), anti-myc (ab9106, Abcam), anti-myc mouse (kindly gifter dy Dr. I. Deyev (Shtykova et al. 2019)).

### DNA methylation analyses

Genomic DNA was extracted with DNeasy Blood & Tissue Kits (QIAGEN). DNA was subjected to bisulfite conversion using EZ DNA Methylation Kit (Zymo Research) and amplified using primers corresponding TRIM25 promoter region for 5’-TTGAATTCTTAGATGAGTGTTGGGAAGG Rev 5’-TTGGATCCAATCGAAACACAACTACTACACC. PCR product was used as a DNA template to make Illumina compatible library with NEB E7645S kit according to the manual. The library was sequenced on HiSeq 1500 in SR mode with 250 bp read length. Alternatively, PCR product was cloned to t-vector (pAL2-T, Evrogen, Russia) and sequenced by Sanger for at least 10 clones for each point.

### EMSA

Kaiso without BTB/POZ domain tagged with GST (339-2016) was purified from transformed BL21 E.coli using GST-sepfarose (Glutathione Sepharose 4B, Cytiva). To generate a probe we amplified sequence containing TRIM25 promoter region using biotin-labeled primers TRIM25for1 5’-biotinGGTTGGCCCACAATATAACCAG, TRIM25for2 5’-biotinGGGAGCTCTTGGGGATCGGA, TRIM25for3 5’-biotinTTCAGGGACTGCTCCTCTCGA, TRIM25rev1 5’-AAGCTGACGCCTGGGTGCAG, TRIM25rev2 5’-AAGCCGTCAGGAAGTCACGTG, TRIM25rev3 5’-GAGCACGACAGCTCCTCGGC. Then half of PCR product was methylated using M.SssI methylase (ThermoFisherScientific) and purified using Qiagen kit PCR purification kit. Binding reaction was performed using LightShift EMSA Optimization and Control Kit (20148X) (Thermo Fisher Scientific, USA). DNA-protein complex was loaded to 5% PAAG (0,5X TBE). Resolved complex was detected using Chemiluminescent Nucleic Acid Detection Module (89880) (Thermo Fisher Scientific, USA).

### Immunoprecipitation and co precipitation analyses

Immunoprecipitation was performed with anti-Kaiso 6F, anti-myc mouse, control IgG antibodies and HA-agarose.

## RESULTS

### Kaiso is involved in TRIM25 promoter methylation

Previously, we generated model cell lines based on HEK293 cells: Kaiso deficient cells (Kaiso KO) and with K42R point mutation that prevents Kaiso SUMOylation (fig1a). We demonstrated that Kaiso is involved in regulation of TRIM25 promoter activity. Kaiso deficiency led to TRIM25 upregulation, while deSUMOylated K42R form of Kaiso repressed its activity (S. Zhenilo et al. 2018). Western blot analyses of total cell lysates confirmed that in K42 mutant cells TRIM25 is downregulated and upregulated in Kaiso deficient cells (fig1b). In order to determine whether the change in TRIM25 transcription is associated with a change in the methylation level of its promoter, we carried out bisulfite analysis. Genomic DNA from wild type cells, Kaiso deficient cells, K42R mutant cells were converted by bisulphate. After the bisulfite conversion region corresponding to the TRIM25 promoter was amplified and analyzed by NGS sequencing. In wild type HEK293 cells level of TRIM25 promoter methylation was about 40% (fig.1c). Kaiso deficiency results in slightly decreased methylation to 30%. This data was confirmed by PCR products cloning into t-vector following Sanger sequencing (Suppl fig1). Reduction of TRIM25 promoter methylation was in accordance with increased transcription of TRIM25 in Kaiso KO cells. This reduction was reverted by expression of exogenous Kaiso (fig1c).

**Figure 1.**
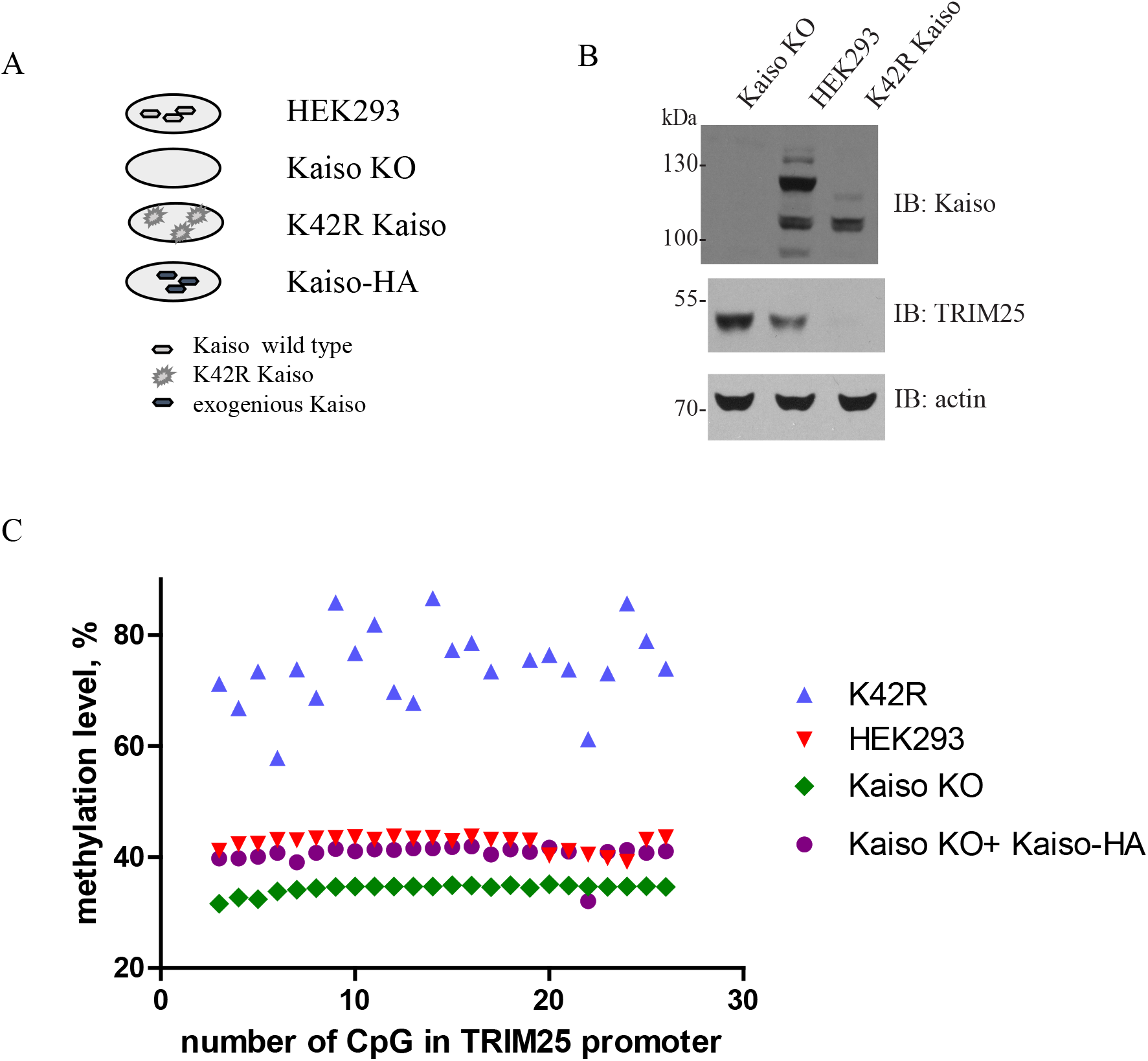
Kaiso regulates TRIM25 promoter methylation. A, scheme of cell lines used in this work. B, western blot analyses of cell lysates from HEK293 cells, Kaiso deficient and K42R mutant cells for expression of TRIM25(S. Zhenilo et al. 2018). C, analysis of promoter TRIM25 methylation by bisulfite conversion and NGS sequencing.

In K42R cells with deSUMOylated Kaiso we detected increased TRIM25 promoter methylation up to 80%. Increased methylation correlates with repression of TRIM25 transcription. Consequently, a deSUMOylated form of Kaiso leads to TRIM25 promoter hypermethylation and may be involved in DNA methylation establishment.

### Kaiso interacts with the methylated promoter of TRIM25

Early, we demonstrated via chromatin immunoprecipitation that Kaiso was detected on TRIM25 promoter in HEK293 cells and in K42R cells. However, it still remains unknown whether Kaiso can directly interact with the TRIM25 promoter region and how this interaction depends on methylation status of DNA. To resolve this question we divided TRIM25 promoter into three fragments. Distal fragment and fragment around TSS contain KBS (CTGCNA) (fig.2). We amplified them with biotin labeled oligonucleotides and methylated obtained PCR products with m.SssI methylase. Despite the fact that two of three probes contain CTGCNA, none of them in the absence of methylation did not interact with Kaiso as it was shown by electrophoretic mobility shift assay. While their methylation resulted in Kaiso’s binding. So, Kaiso binds the methylated promoter region of TRIM25 directly.

**Figure 2.**
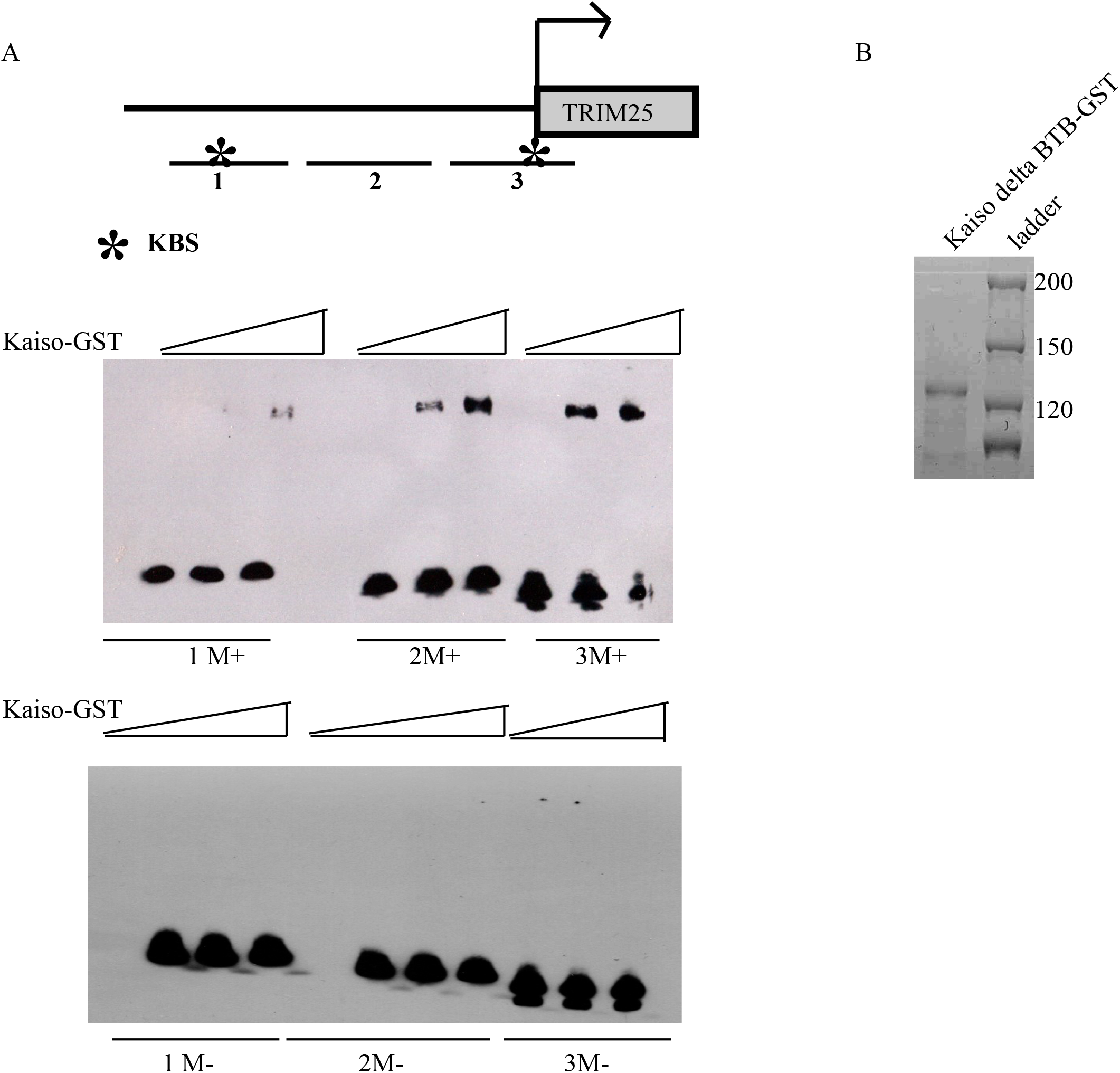
Kaiso directly interacts with methylated TRIM25 promoter. A, EMSA was performed with Kaiso-GST without BTB/POZ domain with three sequences from TRIM25 promoter in dependence from methylation status. B, Coomassie gel staining of GST-sepharose with Kaiso’s domains fused with GST.

### Kaiso is formed complex with de novo DNA methyltransferases

Since Kaiso can increase the methylation level of TRIM25 promoter, so we assumed that Kaiso may attract de novo methyltransferases DNMT3a or DNMT3b. To check whether Kaiso may form a complex with de novo DNA methyltransferases, we cotransfected Kaiso-GFP and myc tagged DNMT3b in HEK293 cells. Co-immunoprecipitation with myc antibodies revealed that Kaiso forms a complex with DNMT3b (fig.3a). To confirm this complex formation we performed immunoprecipitation with antibodies against Kaiso. Western blot analyses demonstrated that Kaiso forms a complex with DNMT3b. Further we want to determine which domains of Kaiso and DNMT3 methyltransferase are essential for complex formation. Kaiso consists of BTB/POZ domain and C-termini zinc finger domain. BTB/POZ domain is responsible for protein-protein interaction. So, we investigated how important is the BTB/POZ domain of Kaiso for interaction with DNMT3b. We cotransfected BTB domain tagged with HA along with DNMT3b-myc and immunoprecipitated proteins with myc antibodies. Western blot analyses showed that the BTB/POZ domain of Kaiso is sufficient for complex formation with de novo DNA methyltransferase DNMT3b (fig3b).

**Figure 3.**
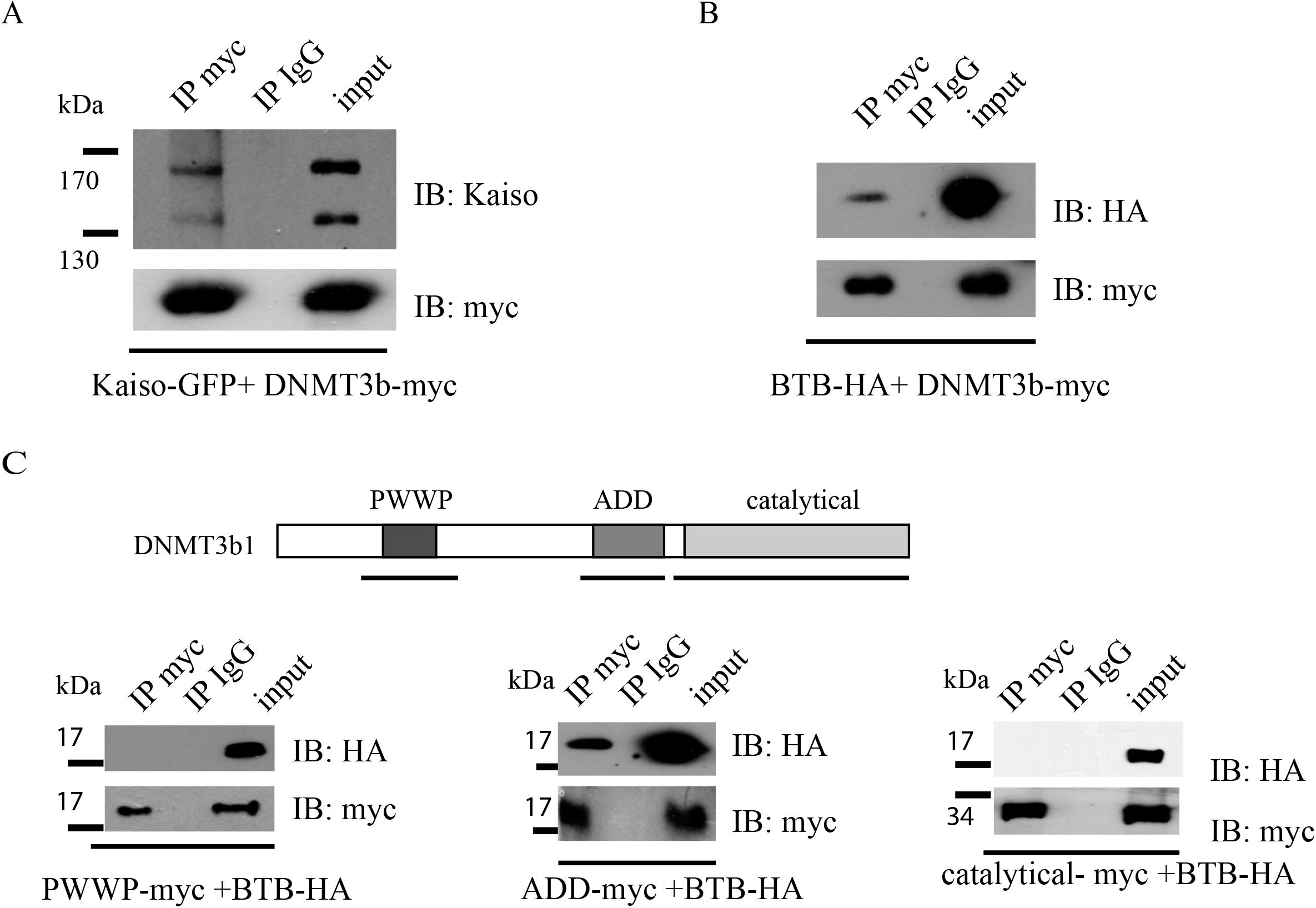
Kaiso and DNMT3b are part of one complex.Kaiso-GFP and DNMT3b-myc (a), BTB-HA and DNMT3b-myc (b), ADD-myc, PWWP-myc, catalitical-myc and BTB-HA (c) were cotransfected in HEK293 cells. Immunoprecipitation was performed with myc and IgG antibodies.

De novo methyltransferase DNMT3b composed of three domains: catalytic domain and chromatin readers domains, ADD and PWWP. We subcloned these domains with myc tag and cotransfected them with BTB-HA. Immunoprecipitation with myc antibodies showed that ADD domain is involved in complex formation with the BTB domain of Kaiso (fig3c). Next, we investigated whether de novo DNA methyltransferase directly interacts with Kaiso or they just formed a multisubunit complex with each other. To resolve this question we performed co precipitation assay. Full length Kaiso, its BTB/POZ domain, spacer region and zinc finger domain were tagged with GST and produced in the *E.coli* expression system. We performed pull down with total cell lysates from HEK293 transfected with DNMT3b-myc. Western blot analyses of precipitated proteins revealed that DNMT3b and Kaiso as well as its domains can not interact directly with each other (data not shown). Consequently, they can exist in one complex, but they do not interact directly.

## Discussion

Establishment of DNA methylation is a complicated process that depends on many factors. Usually as soon as DNA methylation was established after embryo implantation it remains stable except cellular differentiation or some external influence or disease progression. But DNA methylation is flexible in nervous cells that are involved in memory formation, behavioral regulation, learning ability (Lister and Mukamel 2015; Guerrero et al. 2020; Yu, Baek, and Kaang 2011). Methyl DNA binding proteins may be divided into two classes. One class is essential for organism survival. Factors from this class are involved in the establishment of DNA methylation and repression on various repeating elements, regulation of imprinted loci, for example Zfp57 KRAB protein. Also, proteins from this class are important for DNA methylation maintenance such as UHRF proteins. But there exists another class of methyl DNA binding proteins that is highly important for Xenopus laevis or danio rerio development, but loses its significance in mammalian development. These are MBD1, MBD2, MeCP2, Kaiso proteins. Deficiency or mutation in these proteins in mice or humans revealed their importance for normal nervous cell functioning, behaviour, memory formation, immune system working, cancer progression. These proteins attract various corepressor complexes with histone deacetylases influencing histone modifications ((Le Guezennec et al. 2006; Villa, Morey, and Raker 2006; Nan et al. 1998; Yoon et al. 2003)).

In the present work we demonstrated that Kaiso is involved not only in the establishment of repressive histone modifications, but also in the establishment of proper DNA methylation patterns. In HEK293 cells with deSUMOylated Kaiso we observed hypermethylation of TRIM25 promoter along with repression of TRIM25 transcription.While cells deficient for Kaiso showed a decrease in methylation level of TRIM25 promoter and we detected higher level of TRIM25 expression. So, we hypothesized that Kaiso may be involved in de novo DNA methylation followed by heterochromatin formation. This assumption was confirmed by coimmunoprecipitation analyses. Kaiso was detected in one complex with DNMT3b de novo DNA methyltransferase. However, directly they did not interact.

Thus, we can conclude that Kaiso may form a complex with de novo methyltransferase DNMT3b and this may be a novel mechanism of DNA methylation establishment. In what parts of the genome it is important and in what type of cells will be investigated in the future.

## Acknowledgments

We thank Dr. Albert Reynolds for giving anti-Kaiso antibodies. We thank Dr I.Deyev for giving anti-myc mouse antibodies.

